# High-content Imaging-based Pooled CRISPR Screens in Mammalian Cells

**DOI:** 10.1101/2020.06.30.179648

**Authors:** Xiaowei Yan, Nico Stuurman, Susana A. Ribeiro, Marvin E. Tanenbaum, Max A. Horlbeck, Christina R. Liem, Marco Jost, Jonathan S. Weissman, Ronald D. Vale

## Abstract

CRISPR (clustered regularly interspaced short palindromic repeats) -based gene inactivation provides a powerful means of linking genes to particular cellular phenotypes. CRISPR-based screening has mostly relied upon using large genomic pools of single guide RNAs (sgRNAs). However, this approach is limited to phenotypes that can be enriched by chemical selection or FACS sorting. Here, we developed a microscopy-based approach, which we name optical enrichment, to computationally select cells displaying a particular CRISPR-induced phenotype, mark them by photo-conversion of an expressed photo-activatable fluorescent protein, and then isolate the fluorescent cells using fluorescence-activated cell sorting (FACS). A plugin was developed for the open source software μManager to automate the phenotypic identification and photo-conversion of cells, allowing ~1.5 million individual cells to be screened in 8 hr. We used this approach to screen 6092 sgRNAs targeting 544 genes for their effects on nuclear size regulation and identified 14 bona fide hits. These results present a highly scalable approach to facilitate imaging-based pooled CRISPR screens.

## INTRODUCTION

High throughput sequencing in combination with CRISPR technology has greatly accelerated discoveries in biology through unbiased identification of many new molecular players in key biological processes^1–4^. Using a high diversity single guide RNA (sgRNA) library, large numbers of genes can be manipulated simultaneously in a pooled manner and the sgRNA abundance differences can be determined with high throughput sequencing in a short amount of time with low labor and financial cost. Thus far, pooled CRISPR screens have been limited to phenotypes that can be transformed into sgRNA abundance differences, including growth effects^5–7^ or phenotypes that can be directly examined by flow cytometry^8^ or single cell molecular profiling^9–14^. However, many important cellular phenotypes can only be detected under a microscope, which requires a robust method for transforming optically identified phenotypes into differences in sgRNA abundance. Recently, several *in situ* sequencing^15,16^ and cell isolation methods^17–20^ were developed which allow microscopes to be used for screening. However, these methods contain non-high throughput steps that limit their scalability.

Here, we report an imaging-based pooled CRISPR screening method using optical enrichment by automated photo-conversion of a photo-activatable fluorescent protein. Similar to traditional enrichment based pooled CRISPR screens, cells are infected with a sgRNA library and high throughput sequencing is used to examine sgRNA abundance. Instead of traditional enrichment strategies, we use optical enrichment— cells exhibiting the desired phenotype are identified and photo-activated automatically under a microscope. Photo-activated cells are then isolated using flow cytometry. We evaluated this approach using a synthetic fluorescent reporter to estimate screening accuracy and capacity. We then applied this approach to identify genes that regulate nuclear size. This approach is modular, fast, allows millions of cells to be screened within a few hours, and can be easily scaled up to a genome wide level.

## RESULTS

### An Optical Approach for Cell Enrichment by Patterned Illumination followed by FACS Sorting

We developed an approach, which we term optical enrichment, to select cells of interest using a microscope and mark them by photo-conversion, enabling cell isolation using FACS (Fig. 1a). To achieve this, we engineered hTERT-RPE1 cells expressing the photo-activatable fluorescent protein PA-mCherry and observed them under a microscope. Cells of interest were selected by automated image analysis and then photo-converted with patterned illumination using a digital micromirror device (DMD) (Fig. S1a). To avoid undesired photo-conversion of neighboring cells, we limited the conversion pattern to nuclei as identified by the H2B-mGFP signal (Fig. S1b). We developed a plugin for the open source microscope control software μManager^21^ called Auto-PhotoConverter that automates these steps and has a pluggable interface for image analysis code so that it can be used for any desired phenotype (https://github.com/nicost/mnfinder) (Fig. S1c). After harvesting the cells, the photo-converted cells were isolated by FACS. By varying the conversion time of the PA-mCherry, we were also able to create different populations of cells of different intensities, that were clearly distinguished by FACS (Fig. 1b and 1c). This approach makes it possible to analyze multiple phenotypes simultaneously, as discussed below.

**Fig. 1|.**
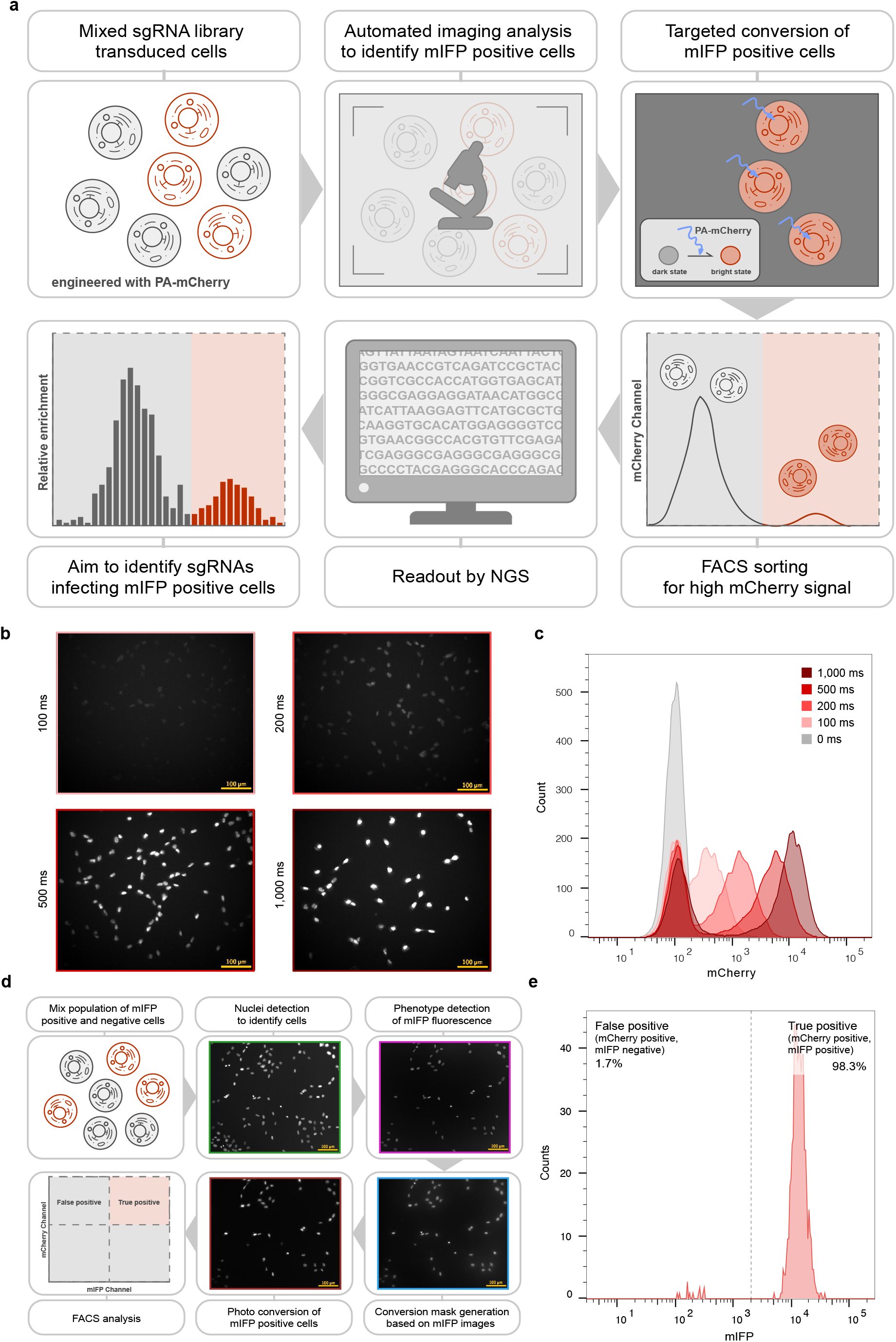
Imaging based Pooled CRISPR Screen. **a**, Schematic of imaging based pooled CRISPR screen. Cells expressing PA-mCherry are infected with pooled sgRNA library and imaged using a microscope. Images are collected and analyzed automatically to generate a conversion mask targeting cells of interest. Exposure with blue light photo-converts cells of interest into mCherry positive cells that are subsequently isolated by FACS. Samples are analyzed by high-throughput sequencing for sgRNA identification. **b-c**, hTERT-RPE1 PA-mCherry cells can be converted into different fluorescent intensity levels **b** that are clearly distinguished by FACS **c**. 100ms is enough for successful photo-conversion. Scale bar: 100μm. **d**, Schematic of experiment to measure specificity. mIFP expression was used as a phenotype to measure specificity. mIFP positive (hTERT-RPE1 PA-mCherry H2B-mGFP mIFP-NLS) and mIFP negative cells (hTERT-RPE1 PA-mCherry H2B-mGFP) were mixed and analyzed under GFP and mIFP channel separately. A conversion mask was generated for each mIFP positive cell. Cells identified by FACS to be mIFP and mCherry double positive are true positives while mCherry positive cells without mIFP fluorescence result from accidental conversion (false positive cells). Specificity is defined as the percentage of true positives of all mCherry positive cells. Example images of image analysis (GFP channel, —; mIFP channel, —), photo-conversion (Blue light channel, —), after photo-conversion (mCherry channel, —) are shown. Scale bar: 100μm. **e**, Cells of interest can be successfully isolated with high accuracy. Example FACS data is shown. Specificity measured at different percentage of mIFP positive cells in the initial mixture is shown in **Fig. S2**.

We next tested the specificity of the photo-conversion approach using a mixture of cells expressing the fluorescent marker mIFP and cells not expressing mIFP. The Auto-PhotoConverter plugin was used to identify and generate a conversion mask based on mIFP fluorescence, the mIFP expressing cells were photo-converted (yielding mIFP-mCherry double positive cells). All cells were then collected and analyzed by FACS (Fig. 1d). We calculated the specificity of this assay from the ratio of true positive and all positive cells (number of cells positive for both mCherry and mIFP divided by all mCherry positives x100%). When the initial subset of mIFP positive cells was 30%, the specificity was calculated as 98.3% (Fig. 1e). The specificity varied with the percentage of mIFP positive cells at the beginning of the experiment and ranged from 80% to ~100% (initial percentage of mIFP positive cells ranges in between 0% and 50%) (Fig. S2). These results indicate the assay yields reliable hit identification regardless of the percentage of hits in the library.

### Optical enrichment enables accurate sgRNA Identification

Having established that we can recover photo-converted cells with high specificity, we next tested if we can successfully identify specific sgRNAs sequences present in these cells. For these experiments, we again selected cells expressing mIFP. Control cells and mIFP positive cells were separately infected with two different sgRNA libraries at a low multiplicity of infection (MOI) to guarantee single sgRNA per cell (Fig. 2a). These two populations were then mixed at a ratio of 9:1 mIFP negative: mIFP positive. A total of 6843 sgRNAs were infected in this mixed population and two biological replicates were performed separately. At least 200-fold coverage of the sgRNA library was guaranteed throughout the screen, including library infection, selection, imaging and FACS. For each replicate, we screened a single imaging plate. A total of 1,825,740 and 1,490,188 RPE-1 cells in the two replicates, were imaged separately using a 20x objective. Automated imaging and photo-conversion of the plate took ~8 hr. The mCherry positive cells and the control population were separately prepared for high throughput sequencing for sgRNA information extraction.

**Fig. 2.**
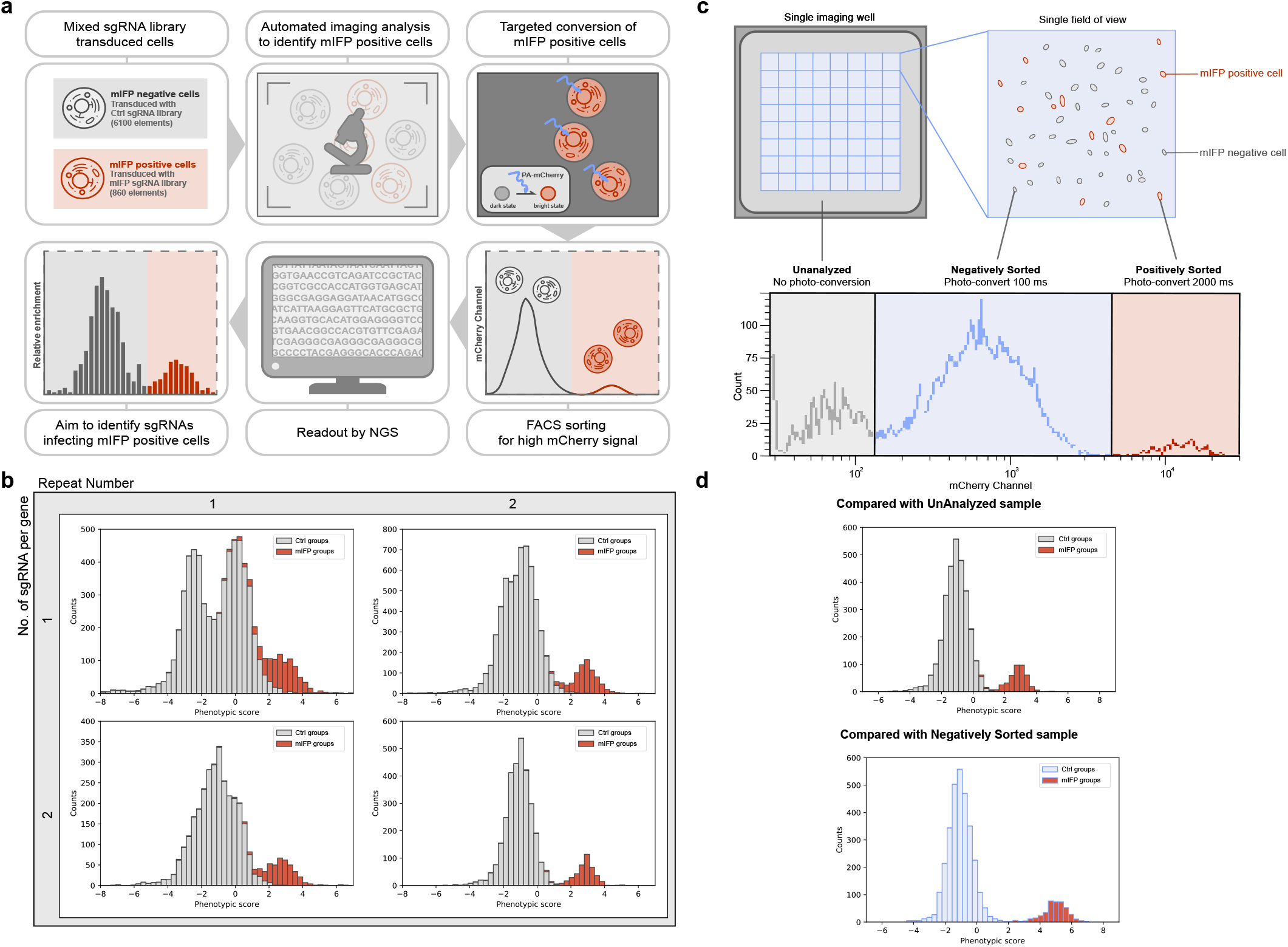
Optical Enrichment enables Accurate sgRNA Identification in a Pooled CRISPR Screen. **a**, Schematic of mIFP proof-of-principle screen. mIFP is used to identify “positive” cells as in **Fig. 1d**. However, mIFP positive and mIFP negative cells were separately transduced with two different sgRNA libraries. Mixed population of mIFP positive and negative cells were then imaged and converted as described in **Fig. 1d**. mCherry positive and mCherry negative cells were then isolated by FACS and prepared for high throughput sequencing to extract sgRNA information. **b**, Distribution of phenotypic scores of all the sgRNAs or grouped sgRNAs in 4 different analysis mode. Top left: single sgRNA/group, single replicate; Top right: single sgRNA/group, average of 2 replicates; Bottom left: 2 sgRNAs/group, single replicate; Bottom right: 2 sgRNAs/group, average of 2 replicates; Red: mIFP groups; Grey: control groups. **c**, Schematic of dual-conversion experiment. Image acquisition generally does not cover the complete imaging well which leaves cells not imaged and unanalyzed. Experiment as described in **Fig. 2a** but mIFP negative cells were also photo-converted (100 ms). mIFP positive cells were converted using a longer conversion time (2000 ms) to guarantee a clear distinction by FACS. Lower panel shows an example of FACS data. **d**, Phenotype identification is improved by comparing with true negative cells rather than unanalyzed cells. Distribution of phenotypic scores of all the groups compared with either unanalyzed sample or negatively sorted sample. 2 sgRNAs were randomly grouped. Data is averaged between 2 replicates. Red: mIFP groups; Grey or Blue: control groups.

For simplicity, we use the terms “mIFP sgRNAs” for the sgRNAs used to infect mIFP positive cells and “control sgRNAs” for the sgRNAs used to infect mIFP negative cells. Since the mIFP positive phenotype is not induced by our sgRNA library, we can group different numbers of sgRNAs for analysis (*i.e.* 1 sgRNA/group or 2sgRNAs/group). These computationally generated groups are used as a proxy for genes in normal sgRNA libraries, as they are also targeted with multiple different sgRNAs. Following sequencing, a phenotypic score was calculated for each sgRNA as its relative enrichment in the mCherry sorted population compared with the control sample. Phenotypic scores of sgRNAs assigning in the same group were then averaged to give a phenotypic score of the given group (Supplementary file 1).

Our results show that, mIFP sgRNAs could be distinguished from control sgRNAs in a single experimental replicate (Fig. 2b, top left). Combining data from both replicates significantly improved segregation of the mIFP and control groups (Fig. 2b, top right). Not surprisingly, the greater the number of sgRNAs assigned to a group (see above), the better the detection of sgRNA hits (Fig. 2b, bottom). Two sgRNAs per group is enough for a reliable screening result, even using a single replicate (Fig. 2b bottom left). Thus, we demonstrate that pooled CRISPR libraries can be screened for phenotypes under a microscope by optical enrichment.

### Improved Phenotype Identification through Multi-parameter Labeling

In most pooled CRISPR screens, only cells showing the phenotype of interest are selected and the relative enrichment of a given sgRNA is calculated based on comparison with the whole cell population. However, this whole cell population is usually collected separately and includes both positive and negative cells, which reduces the perceived enrichment in the positive population (Supplementary file 1). We therefore investigated calculating the relative enrichment of a given sgRNA by comparing with the true negative cells. Not all mCherry negative cells are true negative cells since there are unanalyzed regions outside of the microscope field of view (grey region in Fig. 2c top panel) and cells that fail to pass the filters for phenotype identification (Supplementary file 2). Thus, true negative cells also need to be labeled before harvesting. This task requires selecting for multiple phenotypes simultaneously. We achieved this within the same experiment using different photo-conversion times for true positives (2 sec) and true negatives (100 ms) and then separating them by FACS (Fig. 2c). For comparison, cells going through the same experimental procedure but were not analyzed during image analysis (unanalyzed cells, mCherry negative cells) were also collected to determine the sgRNA composition in the total cell population. As shown in Fig. 2d, the peaks indicating mIFP genes and control genes were separated to a much greater extent when compared with true negative cells than with the whole cell population (unanalyzed sample). This result suggests that this approach indeed can significantly improve screening pooled sgRNA libraries. Additionally, other than labeling true positives and true negatives, this approach can also be used to screen for multiple different phenotypes which greatly expands the phenotypes that could be studied with our approach.

### Pooled CRISPR Screen for Factors involved in Nuclear Size Regulation

To further test our screening method, we performed a screen for regulators of nuclear size. We generated a CRISPRi library of 6092 sgRNAs representing 544 genes whose translation efficiency is upregulated during the G2 phase of the cell cycle. This library includes sgRNAs targeting FBXO5, which is known to cause larger nuclei after knock down^22,23^, and served as the positive control. For this experiment, hTERT-RPE1 cells were further engineered with CRISPRi modality (dCas9-KRAB) to inhibit transcription of genes targeted by the sgRNA library. We selected 2 control sgRNAs that have no targeting sites in the human genome, and as expected had no discernible effect on nuclear size (Fig. S3). Nuclear sizes were measured for control cells and the value of the top 0.5% was used as the screening threshold. As shown in Fig. 3a, H2B-GFP fluorescence of cells infected with this sgRNA library was imaged using an epi-fluorescence microscope and nuclear size was determined by automated image analysis (Supplementary file 2).

**Fig. 3.**
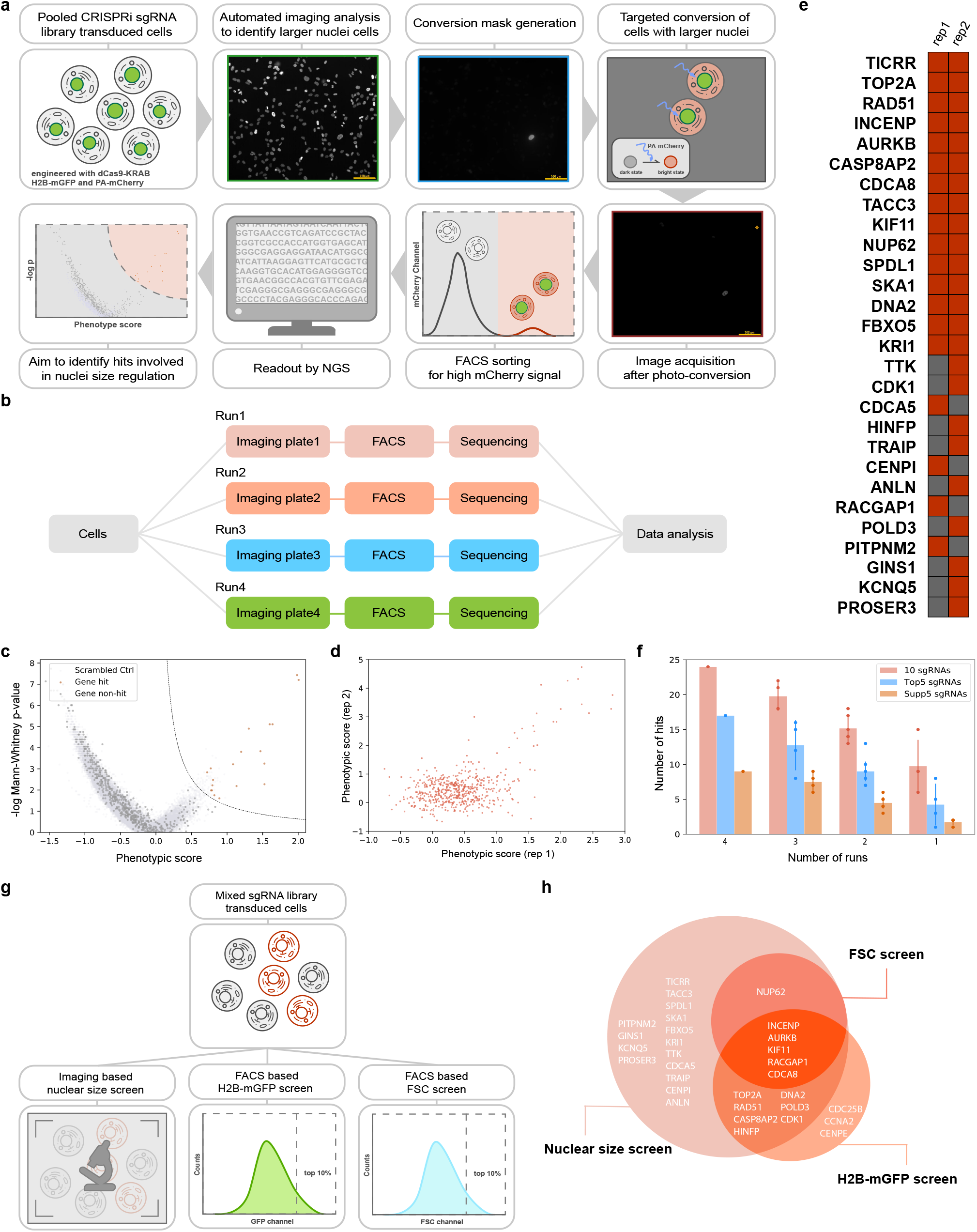
Screens for nuclear size regulators. **a**, Schematic of nuclear size screen. Cells (hTERT-RPE1 dCas9-KRAB PA-mCherry H2B-mGFP) were transduced with a CRISPRi sgRNA library and imaged under the GFP channel. Cells with nuclei larger than 1000 μm^2^ were photo-converted, sorted and analyzed by deep sequencing. Example images of nuclei, (GFP channel, —), photo-conversion (Blue light channel, —), after photo-conversion (mCherry channel, —) are shown. Scale bar: 100 μm. **b**, Work flow of one replicate of the nuclear size screen. For each replicate, transduced cells were seeded into 4 glass-bottom imaging plates. Cells in each single imaging plate were imaged, analyzed, photo-converted, sorted and sequenced separately, termed as separate runs. There were 4 runs in each replicate and sequencing data from these 4 runs was either analyzed separately, or combined (see **f**). **c**, Screening result of one replicate. Detailed calculation was described in **Supplementary file 1** and the other replicate is shown in **Fig. S4**. **d**, Comparison between two replicates. **e**, List of genes identified from two replicates. Red: hit. Grey: non-hit. **f**, Number of hits identified using data averaging from different numbers of runs and/or different library compositions. Error bar: standard deviation. **g**, Work flow of three screens, namely nuclear size screen, H2B-mGFP screen and FSC screen. After transducing the sgRNA library, cells were split and prepared for either imaging analysis (nuclear size screen) or FACS sorting (H2B-mGFP screen and FSC screen). **h**, Comparison of hits identified in FSC screen, GFP screen and nuclear size screen.

Positive cells were photo-activated and sorted together with mCherry negative cells (unanalyzed cells) as a comparison. Two biological replicates were performed, each consisting of 4 imaging plates, containing 5,521,518 and 5,795,313 RPE-1 cells in total respectively. Both replicates were completed within 2 days (each plate taking 7-10 hr). The 4 imaging plates per replicate were carried out as separate screening experiments, termed runs, and data was only combined after sgRNA abundance determination (Fig. 3b). Control groups were generated computationally by randomly regrouping all the sgRNAs and a phenotypic score was calculated for each gene and control group. Data of the 4 runs were averaged and a score summarizing effects from both severity of the phenotype (phenotypic score) as well as confidence level (−log(p value)) was calculated (Supplementary file 1). A value of the top 0.1 percentile of control groups was used as a cutoff for hits (Fig 3c and Fig. S4). The two replicates combined yielded 28 hits of which 15 genes were found in both replicates, including the positive control FBXO5 (Fig. 3d and 3e). To validate the 15 genes that emerged in both replicates of the microscope-based screen for enlarged nuclei, each gene was individually targeted. 14 out of 15 hits (the exception was TACC3) were confirmed to be real hits, with cells exhibiting larger nuclei after knock down (Fig. 4a and Fig. S5).

**Fig. 4.**
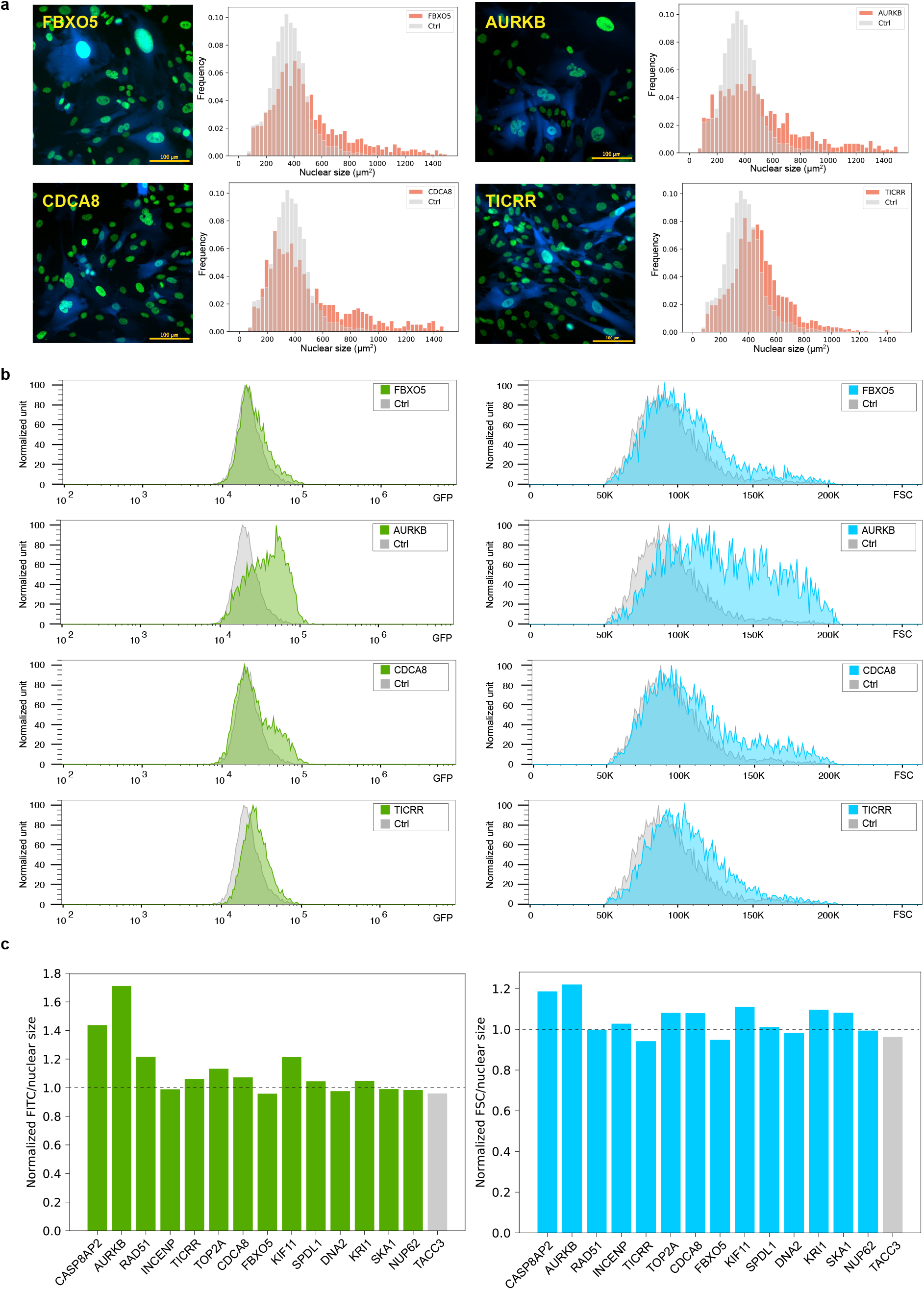
Characterization of Hits Identified in Nuclear Size Screen. **a**, Each hit identified in both replicates was verified under the microscope after infecting with 3-4 sgRNA constructs targeting the gene. Cells were puromycin selected for 2 days before imaging. BFP was expressed from the same sgRNA construct. Example images of 4 hits and their distribution of nuclear sizes are shown in **a**, all the others are listed in **Fig. S5** (at least 1919 cells were analyzed for each gene). **b**, Most cells developed a larger cellular size and higher H2B-mGFP level after knock down. Cells were infected with the same 3-4 sgRNAs/gene and puromycin selected for 3 days before FACS analysis. Example FACS data of the same 4 hits are shown in **b** and all the others are shown in **Fig. S5**. **c**, Most cells maintained a constant ratio between nuclear size and DNA content or cellular size after knock down. TACC3, confirmed to be a control gene was used for comparison (Grey bar).

To estimate the minimum requirements for performing an optical enrichment pooled CRISPR screens, we computationally analyzed the effect of both library composition and number of runs on the screening results. Utilizing data from replicate 2, we re-ran the analysis, comparing results when fewer sgRNAs per gene and/or fewer runs were included. We binned the sgRNAs based upon three commercially available CRISPRi libraries: 10 sgRNAs/gene and the “Top5” or “Supp5” sub-pools of the sgRNA library^24^. “Top5” and “Supp5” libraries were first designed in Jonathan Weissman’s lab by splitting their original 10 sgRNAs/gene library based on predicted sgRNA activity^24^. As expected, using more sgRNAs and/or runs allowed more hits to be identified (Fig. 3f), and the “Top5” sgRNAs yielded more hits than “Supp5” sgRNAs. However, even when using only the Top5 sgRNAs for two runs or 10 sgRNAs with a single run, around 10 hits were successfully identified (Fig. 3f). Thus, based upon factors such as the time to run a screen and available sgRNAs, fewer sgRNAs/gene and/or less runs might be used in a screen with the expectation of a somewhat reduced identification rate.

Since nuclear size often positively correlates with DNA content and cell size, we also sorted cells based upon H2B-mGFP intensity (as a proxy for DNA content) or forward scattering (FSC) signal (cell size) (Fig. 3g and S6). To compare results directly, these two screens were performed at the same time as our imaging-based nuclear size screen (Fig. 3g). The top 10 percentile of cells based on either GFP fluorescence or FSC signal were separately sorted and prepared for high throughput sequencing. In the H2B-mGFP intensity screen, two replicates identified 10 and 12 hits respectively, with 7 in common, while 6 and 0 were identified in the FSC screen (Fig. S6). Together, a total of 16 genes were captured in the H2B-mGFP and FSC screens (Fig. 3h); 13 out of these 16 genes were also identified through the imaging based nuclear size screen. These data suggest that a direct measurement utilizing a microscope can provide significant improvement in hit yield even for phenotypes that could be indirectly screened with other approaches.

## DISCUSSION

High throughput sequencing has transformed our ability to perform pooled genetic screens on a broad scale. However, applying enrichment-based pooled CRISPR screens to optical based phenotypes has been challenging. In this study, we developed a simple imaging-based pooled CRISPR screening method. Using a photo-activatable fluorescent protein PA-mCherry, cells of interest can be easily labeled through photo-activation and isolated with FACS sorting, which enables sgRNA identification by high throughput sequencing. We have combined this optical enrichment strategy with pooled CRISPR-Cas9 to perform imaging based CRISPR screens. Recently, a similar strategy was independently developed by another group, highlighting the broad applicability and power of this approach toward identifying key regulators of previously intractable phenotypes^19^.

### Advantages and Limitations of Phenotypic Screening by Optical Enrichment

Image processing and microscopic operations are the time limiting steps of most imaging-based genetic screens. Our optical enrichment based pooled screening method is fast and scalable. For example, the image analysis code developed for this study can be run on millisecond time scale per field of view, and cells can be completely separated from the control population on a FACS machine with as little as 100 ms photo-conversion time (Fig. 1c). This process is fast, which makes high speed screening of large amounts of cells possible. In our system, 1.5 million RPE-1 cells can be imaged and photo-converted in 8 hr with a 20x objective. This is significantly faster than the *in situ* methods. For phenotypes as penetrant as mIFP positive phenotype, a library of 6092 sgRNAs with 2 sgRNA/gene can be successfully screened with a single replicate. A genome scale screen of such a phenotype can be executed within 3 days (time of image analysis and photo conversion). Even for more complex phenotypes, such as the nuclear size screen described here, a genomic screen can be finished within 2 weeks using the “Top5” sgRNA library and 3 runs. This time can be even shortened with further optimization such as the use of a microscope with a larger field of view, a lower magnification objective, optimization of imaging analysis algorithms, *etc*.

Optical enrichment screening also is possible to undertake for phenotypic screens with relatively low hit rates. In our nuclear size screen, more than 1.5 millions cells were analyzed during each run, but only 2000-4000 cells were recovered after FACS sorting due to the low hit rate (2.76%). With our optimization of small cell number genomic DNA preparation (methods), we are now able to process sequencing samples from only a few thousand cells.

Our optical-enrichment screening approach also has the advantage of being able to screen multiple phenotypes simultaneously by using different photo-activation times. With PA-mCherry, we show that 4 distinct phenotypes could be potentially sorted (Fig. 1c). We demonstrate this in practice by differential photo-activation of true positive and negative cells to improve screening sensitivity. However, differential time of photo-activation could also be applied to analyze different phenotypes. This approach can be further developed by combining multiple photo-activatable fluorescent proteins in each cell.

Our approach also has some limitations. Phenotypes were identified during image analysis/photo-conversion process, thus the analysis code has to be fast and robust. In our study, fluorescent labeling was used to facilitate phenotype identification and a custom analysis code was generated for each phenotype. In our assay, either phenotype only takes couple hundred milliseconds to identify while this requires developing analysis code for each single phenotype. However, this could be overcome by implementing other image analysis stragedies, including deep learning, which will further benefits laboratories which do not have image processing experts. Additionally, our approach is currently not compatible with fixation assay, thus transient phenotypes might be difficult to capture. However, we expect this to be solvable by further optimizing our screening pipeline to make it possible to prepare sequencing samples after fixation.

### Optical Enrichment compared to Other Methods for Phenotypic Screening

Two other methods have recently been developed to use imaging both for phenotypic screening and decoding to permit sgRNA identification in individual cells *in situ*^15,16^. In both methods, CRISPR sgRNA expression constructs were modified to express both a sgRNA and a barcode. The barcode can be read out either by *in situ* sequencing^15^ or sequential fluorescence *in situ* hybridization (FISH)^16^. Both methods require sgRNA to be re-barcoded necessitating *de-novo* design and library re-synthesis preventing reuse of most existing sgRNA libraries. In addition, cells need to be fixed preventing further cell-based assays of the identified cells. Most importantly, both of these methods cannot easily scale to the whole genome because of barcoding limitations and the long imaging time required.

Another newly published method, similar to ours, also uses high throughput sequencing as an end point assay. Instead of using FACS to enrich cells of interest, this method cultures cells on microcraft arrays (magnetic polystyrene particles designed to capture single clones) to enable cell isolation as separate clones (CRaft-ID)^20^. This method also can use most available sgRNA libraries and is compatible with further live cell studies. However, it is difficult to perform a genome wide screen with CRaft-ID, since it requires single cell isolation during cell culture and thus limits the number of cells that can be screened (6000 colonies/array). In addition, CRaft-ID can not be used to screen for phenotypes that cause defects in monoclonal growth, including essential genes. Our assay, on the other hand, provides an option for genome-wide screens and allows for study of genes essential to growth.

### Genes involved in Nuclear Size Regulation

We applied optical enrichment to a screen for genes involved in nuclear size determination. Of the 15 genes that were identified in replicate screens, all have known roles during cell cycle regulation except KRI1 which is involved in cell death regulation in *C. elegans*^25^ (Supplementary file 3). Six genes are involved in spindle function and chromosome segregation, which includes KIF11^26^, NUP62^27^, SPDL1^28^ and three core chromosomal passenger complex (CPC) components INCENP, AURKB and CDCA8^29,30^. Three genes function in DNA damage and repair, namely TICRR^31,32^, TOP2A^33,34^ and RAD51^35,36^, while the remaining four play roles in histone synthesis (CASP8AP2^37^), DNA maintenance (DNA2^38,39^) and cell cycle regulation (SKA1^40,41^ and FBXO5^22,23^) (Supplementary file 3). Some of these functions might directly explain the larger nuclei phenotype after knock down. For example, the loss of FBXO5 was suggested to lead to cellular senescence, resulting in larger nuclei^23^. Knockdown of CPC components including AURKB, INCENP and CDCA8 leads to asymmetrical distribution of nuclear material and cytokinesis failure, thereby generating abnormally large nuclei^29,30^. This notion also matches with the observation that FBXO5 knock down produces only larger nuclei while knock down of CPC components also leads to smaller nuclei, especially at later stages (Fig. S7).

To begin to understand the mechanism underlying nuclear size regulation of our 14 hits, we investigated changes in DNA content, measured by H2B-mGFP, and cell size, assessed using forward scattering on FACS, after knock down. Almost all hit genes show increases in both H2B-mGFP fluorescence and FSC signal (Fig. 4b and Fig. S5), indicating an unchanged ratio of nuclear size, DNA content and cellular size after knock down (Fig. 4c). This result is expected based upon both the nucleoskeletal theory, which highlights the role of DNA content in nuclear size regulation, and the karyoplasmic ratio theory, which suggests that nuclear size is always related with cellular size^42–45^. Two interesting exceptions were CASP8AP2 and AURKB which have a much higher DNA content/nuclear size ratio (Fig. 4c left panel) that awaits further investigation.

### Conclusion

In summary, our data demonstrate the power of our optical enrichment based pooled CRISPR screening method to study previously inaccessible phenotypes with high efficiency and accuracy. This method is simple, fast, uses open source software, and can be applied to commercial or institutional genome-scale CRISPR sgRNA libraries. A digital micromirror device is required, but this can be introduced into the light path of common commercial microscopes. This screening approach could be broadly applied across many biological phenotypes including morphological changes, sub-cellular organization and cellular dynamics. Pluggable image analysis code enables selection of any desired morphological phenotypes as long as fast and robust detection code can be created, which is also an area highly suited for deep learning approaches. We also anticipate that this screening approach can be integrated with other profiling technologies such as single cell sequencing, further expanding its application to other research fields.

## ONLINE METHODS

### Plasmid Sequences

CRISPRi construct (Addgene 85969) and sgRNA parental construct (Addgene 84832) were a kind gift from Jonathan Weissman lab. Other plasmid constructs used in this study are described in Supplementary file 4.

### Cell Line Generation

#### hTERT-RPE1 CRISPRi

All the hTERT-RPE1 cells were grown in DMEM/F-12 media supplemented with 10% FBS and Pen/Strep (complete DMEM/F-12 medium). CRISPRi modality dCas9-KRAB-BFP construct was stably expressed in hTERT-RPE1 cells via lentiviral infection, as described below. BFP positive cells were sorted after 2 days.

#### hTERT-RPE1 CRISPRi PA-mCherry

The photo-convertible cell line was generated starting with hTERT-RPE1 CRISPRi cell line. The PA-mCherry construct was stably expressed in hTERT-RPE1 CRISPRi cells via lentiviral infection as described below. Monoclonal cell lines were grown and screened under the microscope to select clones with successfully integrated PA-mCherry construct. A cell line that showed high fluorescence and homogeneous activation after photo-conversion was chosen to use in this study.

#### hTERT-RPE1 CRISPRi PA-mCherry H2B-mGFP and hTERT-RPE1 CRISPRi PA-mCherry H2B-mGFP mIFP-NLS

H2B-mGFP and mIFP-NLS constructs were sequentially integrated into hTERT-RPE1 CRISPRi PA-mCherry cells via lentiviral infection. GFP positive cells or GFP/mIFP double positive cells were selected by FACS at 2days post-infection.

### sgRNA Sequences

Two negative control sgRNAs were used in this study and their protospacer sequences (the part of the target sequences) are GCTGCATGGGGCGCGAATCA and GTGCACCCGGCTAGGACCGG. sgRNA libraries used in this study were gifts from Jonathan Weissman’s lab. Since the cell line used (hTERT-RPE1 CRISPRi PA-mCherry) for the mIFP proof-of-principle screen has CRISPRi modality, we used two CRISPRa sgRNA libraries for this screen. These two libraries are described in Supplementary files 5 and 6. The CRISPRi sgRNA library used in the nuclear size screen is described in Supplementary file 7. sgRNAs used for hit verification are listed in Supplementary file 8.

### Lentivirus Preparation and Transduction

For CRISPRi modality construct and sgRNA libraries, lentiviral particles were packaged by transfecting HEK293T in a 15 cm cell culture dishes at 70% confluency with 8 μg plasmid, 1 μg PMD2.G, 8 μg dR8.91, 48 μl TransIT-LT1 transfection reagent (Mirus Bio) and 1300ul serum-free Opti-MEM medium. Medium containing lentivirus was collected 72 hr post-transfection and concentrated 10 fold using an Amicon Ultra Centrifugal Unit (MilliporeSigma). For other constructs including PA-mCherry, H2B-mGFP, mIFP-NLS and small scale sgRNA virus preparations, lentiviral particles were packaged by transfecting HEK293T in a 6-well plate at 70% confluency with 1 μg PA-mCherry plasmid, 0.1 μg PMD2.G, 0.9 μg psPAX2, 10 μl TransIT-LT1 transfection reagent (Mirus Bio) and 250 μl serum free Opti-MEM medium. Medium containing lentivirus was collected 72 hr post-transfection and concentration was not needed. 250 μl supernatant was used to transduce a 6-well plate of corresponding cells by spinning infection at 2000rpm for 1 hr. Polybrene infection reagent (Sigma) was used to increase infection efficiency. Medium was replaced with complete DMEM/F-12 medium immediately after spinning infection. Cells were puromycin selected at 5 μg/ml to select for cells successfully receiving the sgRNA (sgRNA construct harbors puromycin resistance cassette). For screening, cells were puromycin selected for 3 days and for hit verification and follow up analysis, cells were selected for 2 days.

### Microscopy

Cells were grown in 96-well glass bottom dishes (Matriplate, Brooks) after puromycin selection. Images were acquired and analyzed by dpi-fluorescence imaging using Nikon Eclipse Ti-E microscope with a 20x 0.75na (Plan Fluor) objective. A digital micromirror device (DMD, DLP LightCrafter 6500 Evaluation Model, Texas Instrument) was positioned behind the back port of the microscope and illuminated using a Sutter HPX-L5UVLambda LED light source (8W output centered around 405 nm) coupled to a (3 or 5) mm liquid light guide with an optical lens. The DMD image was projected into the sample plane using a 100 mm focal length achromatic doublet lens, and a 1x beam “expander” consisting of a pair of 80 mm focal length achromatic lenses, followed by a 450 nm long pass dichroic mirror positioned under the dichroic mirrors used for epi-illumination (Fig. S2). With all pixels of the DMD in the “on” position, we measured ~40 mW in the back focal plane of the objective. During image acquisition, cells were maintained at a constant temperature of 36°C-37°C using a stage top incubator (Tokai Hit). Camera exposure times were usually set to 500ms for GFP channel; 100ms for mCherry channel and 1000 ms for mIFP channel.

### FACS

For the mIFP proof-of-principle and nuclear size screen, cells were trypsinized and sorted using a BD FACSAria III. For hit analysis, cells were analyzed with BD FACSAria II after 2 days’ of puromycin selection. Cells were gated for single cell population and the GFP and FSC levels were analyzed using Flowjo v10.6.2. Detail gating strategy was provided in **Supplementary file 9**.

### Sequencing sample preparation

Sequencing sample was prepared using a protocol from Jonathan Weissman lab (https://weissmanlab.ucsf.edu/CRISPR/IlluminaSequencingSamplePrep.pdf) except that genomic DNA of positively sorted samples was extracted with the Arcturus PicoPure DNA Extraction Kit for small cell number genomic DNA extraction.

### Data analysis

The screening analysis pipeline was based on a pipeline developed in Jonathan Weissman’s lab (https://github.com/mhorlbeck/ScreenProcessing). The formula used in the calculation is listed in Supplementary file 1.

### mIFP proof-of-principle screen and Nuclear size screen

For the mIFP proof-of-principle screen, mIFP positive cells (hTERT-RPE1 CRISPRi PA-mCherry H2B-mGFP mIFP-NLS) and mIFP negative cells (hTERT-RPE1 CRISPRi PA-mCherry H2B-mGFP) were stably transduced with the “mIFP sgRNA library” (CRISPRa library with 860 elements, see Supplementary file 5) and the “control sgRNA library” (CRISPRa library with 6100 elements, see Supplementary file 6) separately. For the nuclear size screen, cells (hTERT-RPE1 CRISPRi PA-mCherry H2B-mGFP) were stably transduced with the “nuclear size library” (CRISPRi library with 6190 elements, see Supplementary file 7). To guarantee that cells receive no more than one sgRNA per cell, BFP was expressed on the same sgRNA construct and cells were analyzed by FACS the day after transduction. The experiment only continues when 10-15% of the cells were BFP positive. These cells were further enriched by puromycin selection (a puromycin resistance gene was expressed from the sgRNA construct) for 3 days to prepare for imaging. Cells were then seeded into 96-well glass bottom imaging dishes (Matriplate, Brooks) and imaged the next day. Either mIFP positive cells or cells passing the nuclear size filter were identified and photo-converted automatically using the Auto-PhotoConverter Micro-Manager plugin. Cells were then harvested by trypsinization and isolated by FACS. Sorted samples were used to prepare sequencing samples.

## ACKNOWLEDGEMENTS

We would like to thank Luke A. Gilbert, Martin Kampmann and Kara McKinley for helpful discussions. This work was supported by the Howard Hughes Medical Institute.

## AUTHOR CONTRIBUTIONS

M.E.T. conceived of the project with input from R.D.V.; N.S. developed the Auto-PhotoConverter plugin; N.S. developed image analysis code with input from X.Y.; M.A.H. designed sgRNA libraries with input from M.E.T.; X.Y., C.R.L. and M.J. cloned the sgRNA libraries; X.Y. and S.A.R. performed the experiments; X.Y. analyzed the data; X.Y. drafted the manuscript; X.Y., N.S. and R.D.V. edited the manuscript with input from M.E.T. and S.A.R.; and all authors read and approved the final article.

---

**Fig. S1.**
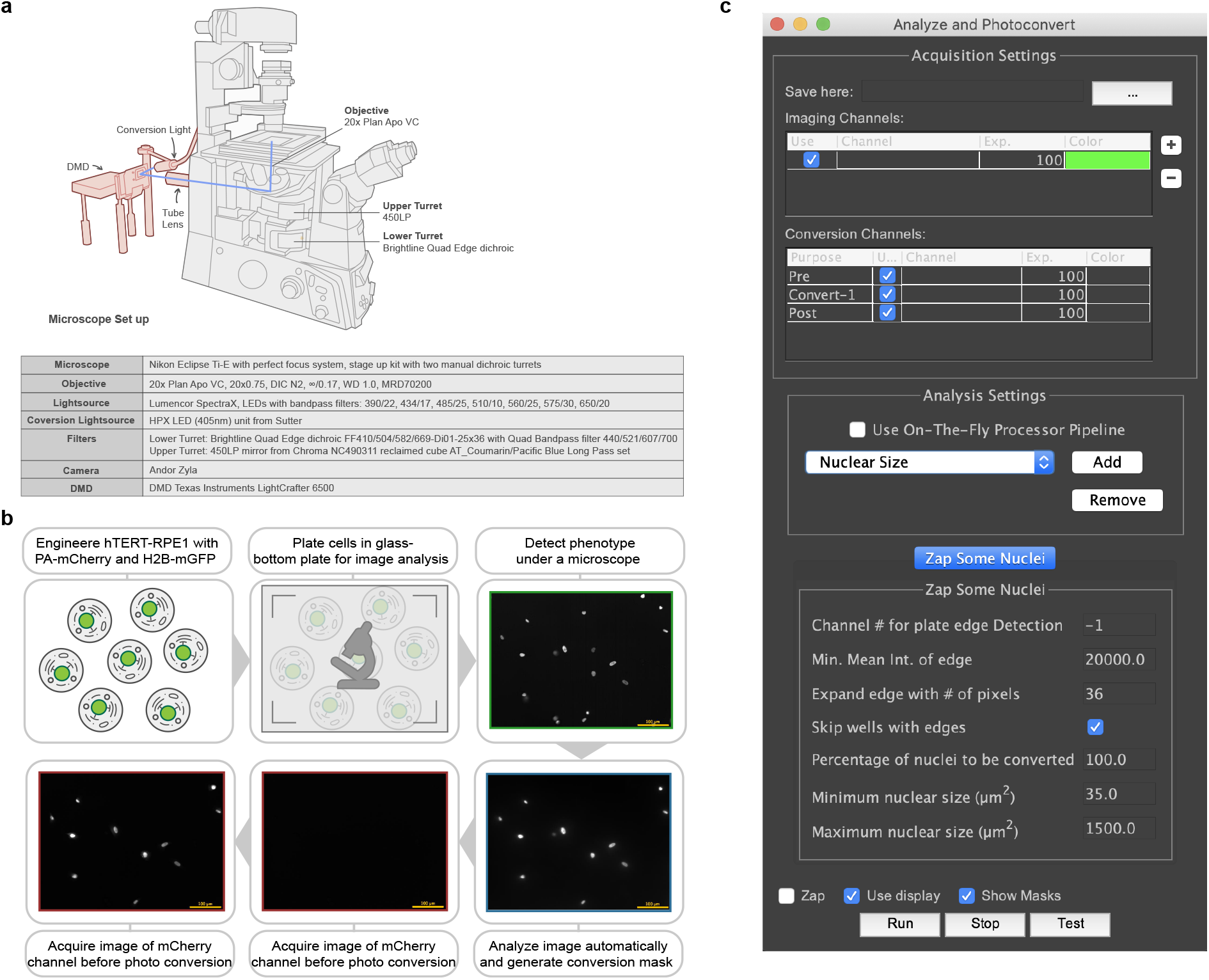
Microscope and μManager Plugin for Photo-Conversion Experiments. **a.** A digital micro mirror device (DMD) and a LED blue (centered around 405nm) light source were engineered on a Nikon Eclipse Ti-E microscope as shown in the figure. A computer was used to control the DMD which reflects light into the microscope only when pixels are in the “on” position, so displaying a mask matching the cell photo-converts that cell. **b**, Schematic of photo-conversion experiment. hTERT-RPE1 PA-mCherry H2B-mGFP cells were seeded in a glass-bottom plate the day before imaging. Example images of cells undergone image analysis (GFP channel, —), photo-conversion (Blue light channel, —), before and after photo-conversion (mCherry channel, —) are shown. Scale bar: 100μm. **c**, A μManager plugin was developed to enable automatic image acquisition, analysis and photo-conversion. An analysis plugin defines its own set of parameters that can be manipulated by the user. Two analysis plugins were used in this study: one for cell identification and another for nuclear size measurement.

**Fig. S2.**
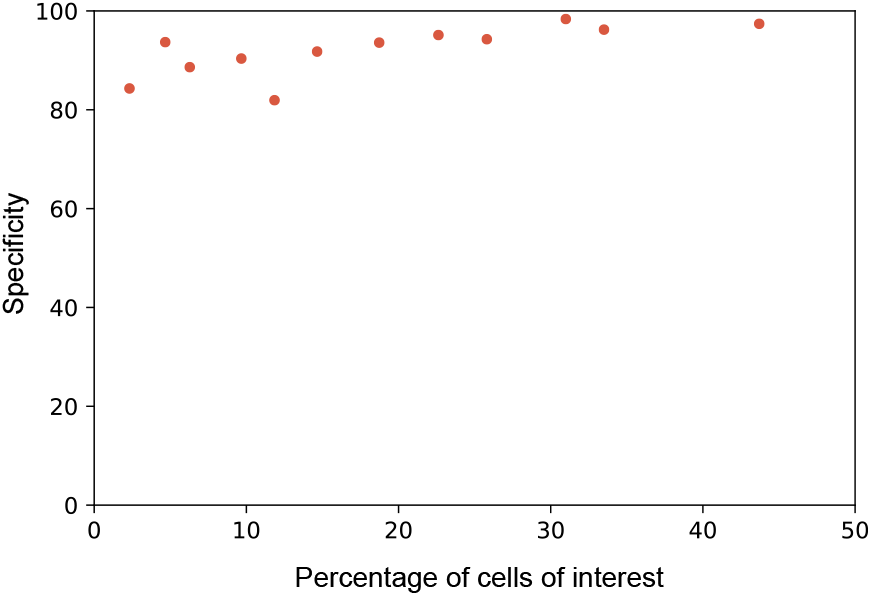
High Photo-conversion Specificity across a Broad Range of Percentage of Hits. Different ratios of mIFP positive cells and mIFP negative cells were mixed to measure specificity at different percentage of hits. mIFP positive cells were photo-converted to become mCherry positive, and FACS analysis was used to measure the fraction of mCherry positive cells that were also mIFP positive (true positives).

**Fig. S3.**
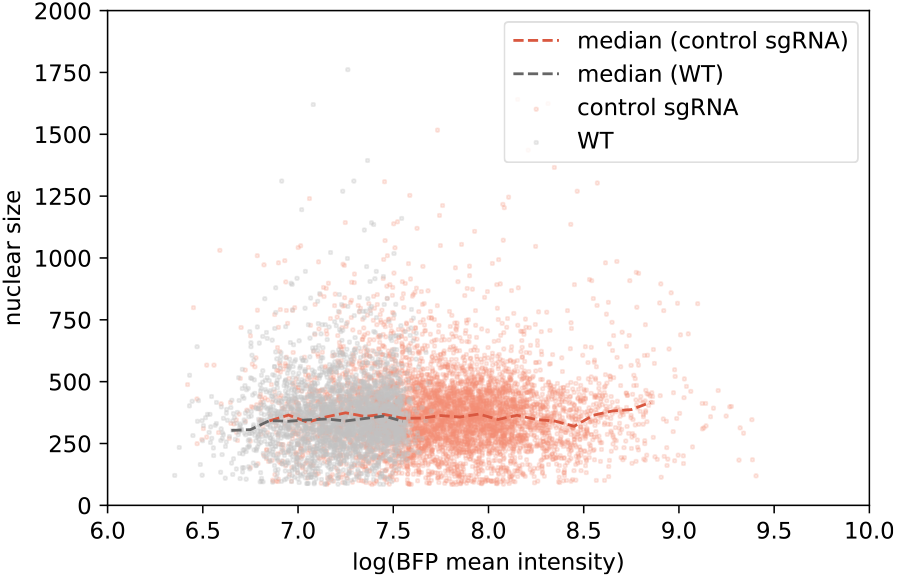
Negative Control sgRNAs do not Affect Nuclear Size after Viral Infection. Two negative control sgRNAs which have no target sites in the human genome were designed. Cells (hTERT-RPE1 dCas9-KRAB PA-mCherry H2B-mGFP) were infected and puromycin selected for 3 days before imaging. A BFP was encoded on the sgRNA construct as a marker for successful infection. FACS results show no correlation between nuclear size and BFP intensity.

**Fig. S4.**
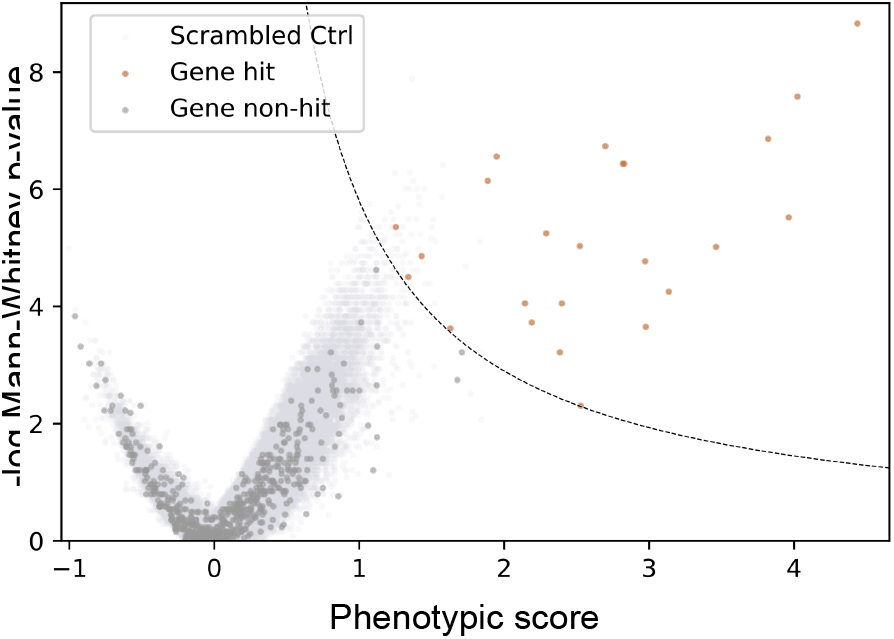
Nuclear Size Screen Result of the Second Replicate. First replicate is shown in Fig. 3c.

**Fig. S5.**
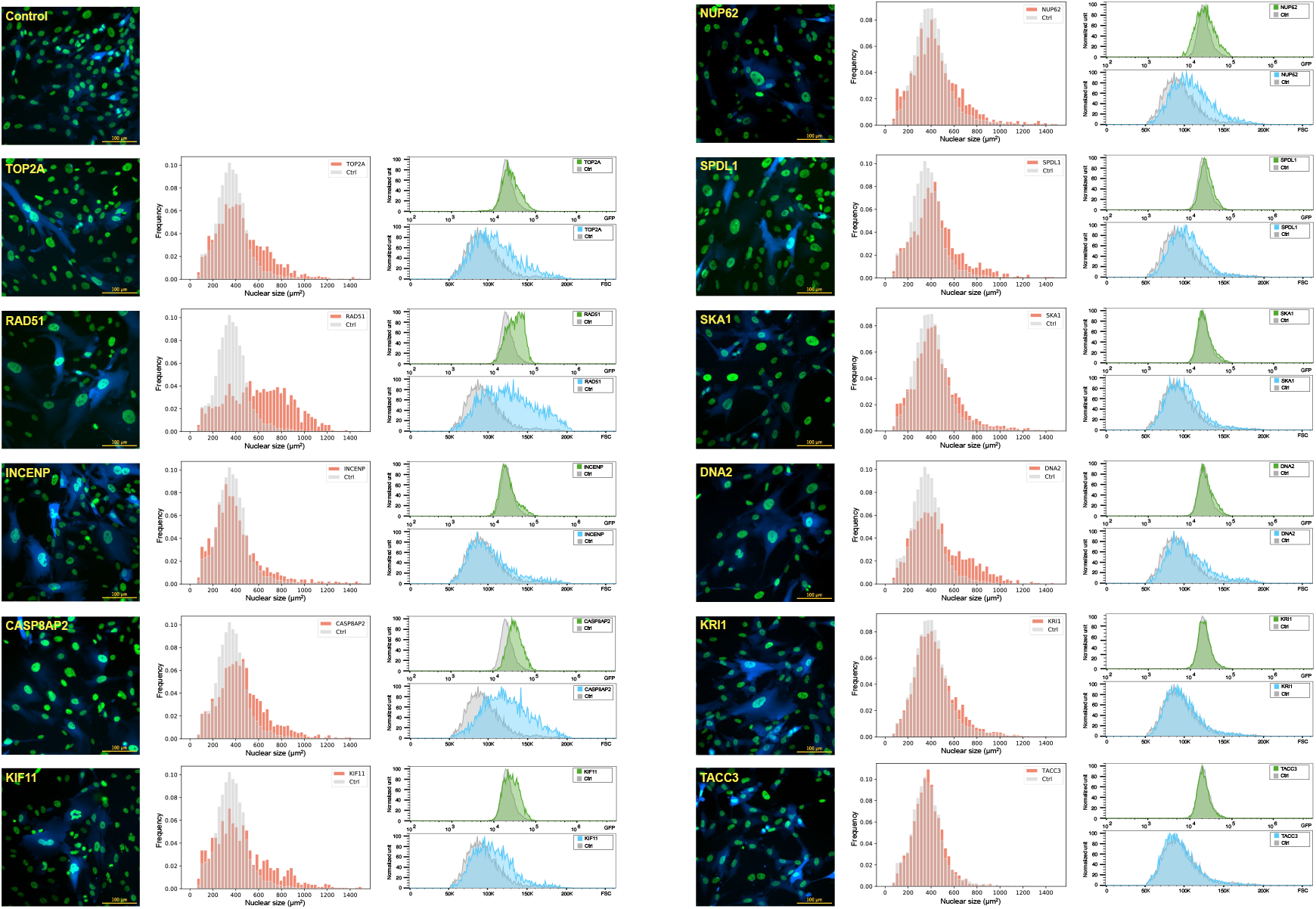
Characterization of Hits Identified in Both Replicates of the Nuclear Size Screens. Example images, distribution of nuclear size, FACS data of H2B-mGFP fluorescence and FSC distribution of each hit (other than the 4 shown in Fig. 4) after knock down are shown (at least 1919 cells were analyzed for each gene).

**Fig. S6.**
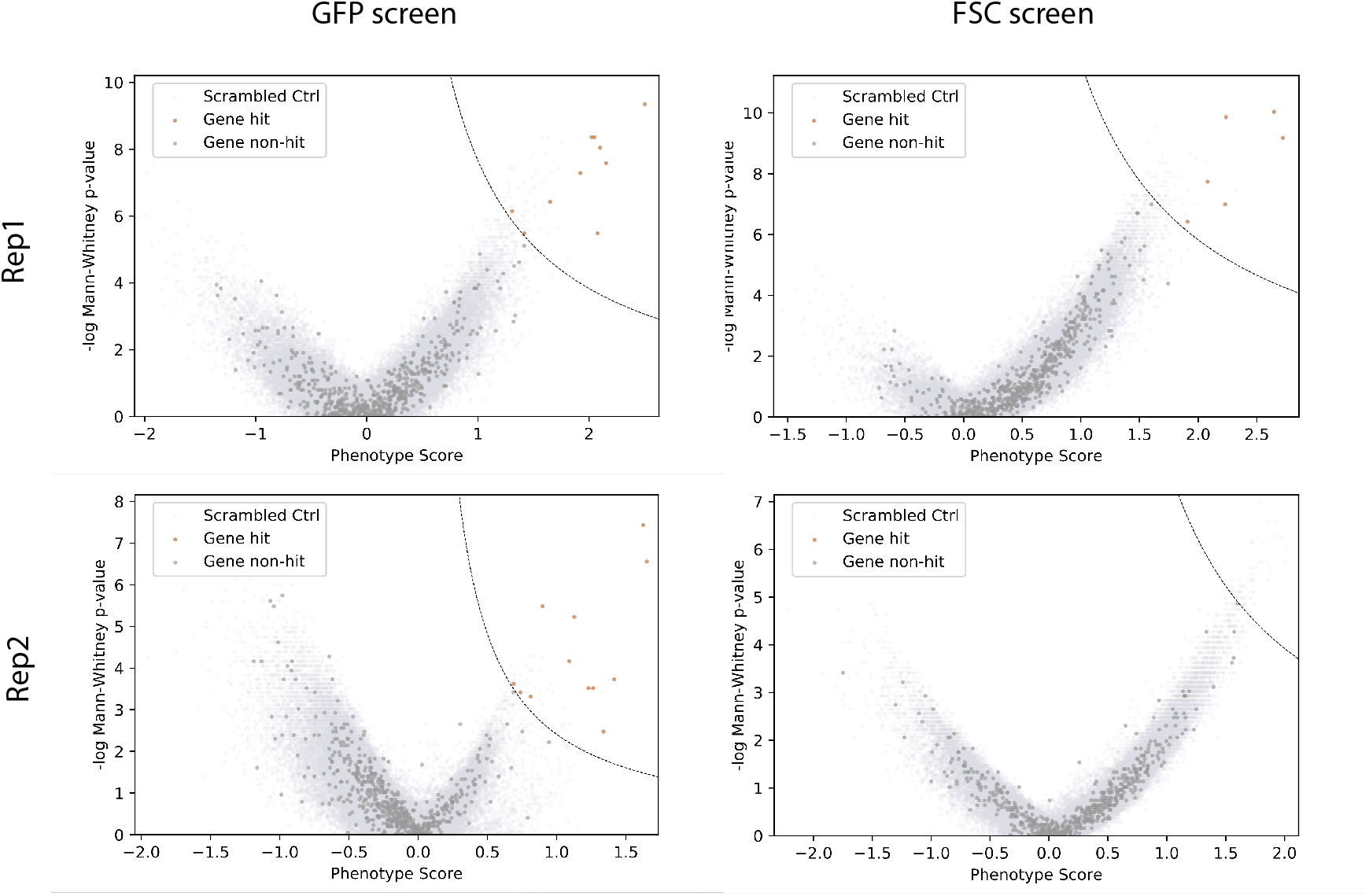
Screen results of FSC and GFP screens. Cells (hTERT-RPE1 dCas9-KRAB PA-mCherry H2B-mGFP) were infected and puromycin selected for 3 days. The top 10 percentile of cells based on either GFP fluorescence or FSC signal were separately sorted and prepared for high throughput sequencing. Screen results of two replicates were shown.

**Fig. S7.**
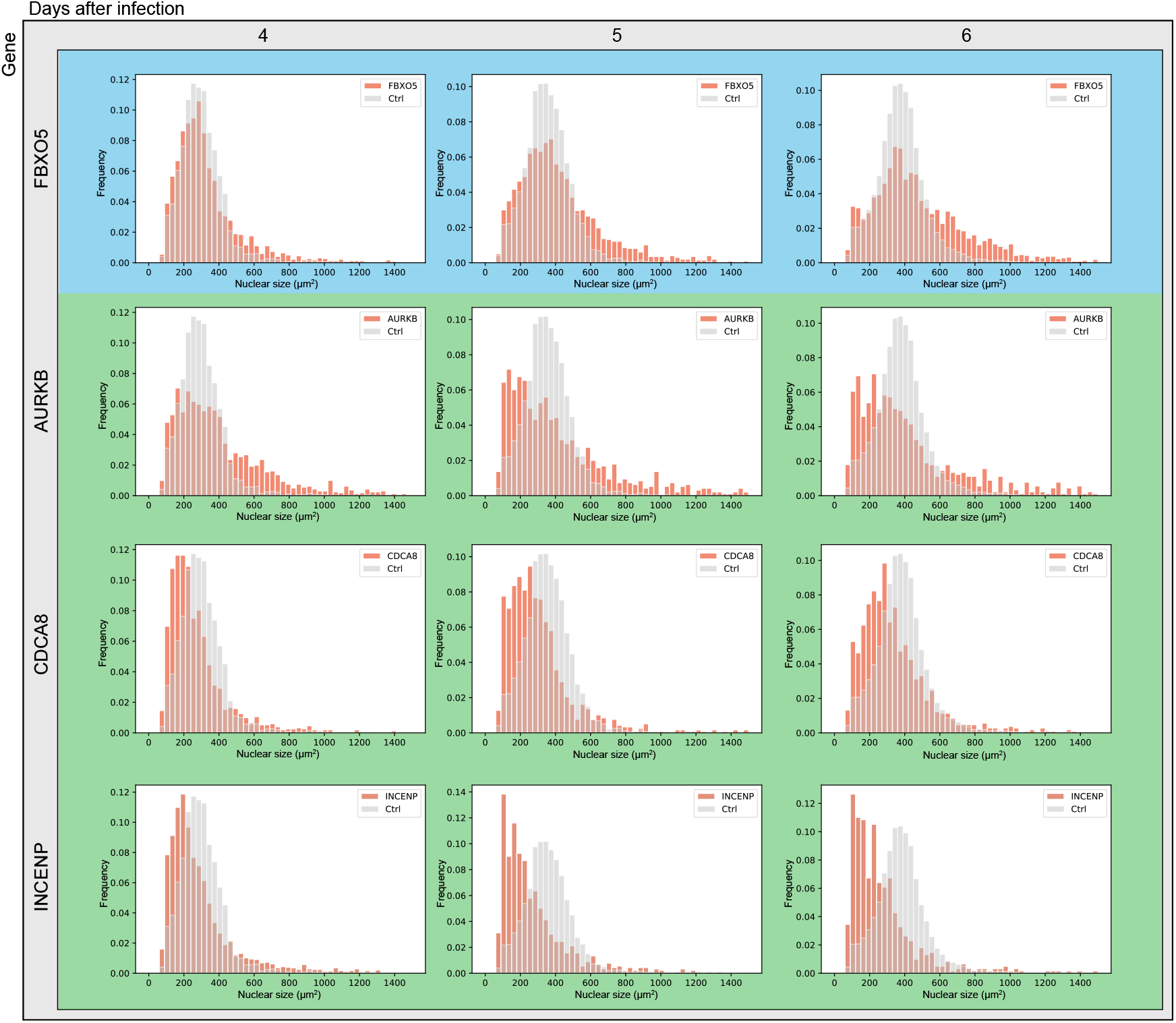
Nuclear Size Distributions Evolve differently after Knock Down. Nuclear size distributions of FBXO5 (blue) and CPC components AURKB, CDCA8 and INCENP (green) were measured by microscopy 4, 5, and 6 days after infection (at least 1163 cells were analyzed).

